# The evolution of sexual dimorphism in gene expression in response to a manipulation of mate competition

**DOI:** 10.1101/2023.08.15.553445

**Authors:** Prashastha Mishra, Howard D. Rundle, Aneil F. Agrawal

## Abstract

Many genes are differentially expressed between males and females and patterns of sex-biased gene expression (SBGE) vary among species. Some of this variation is thought to have evolved in response to differences in mate competition among species that causes varying patterns of sex-specific selection. We used experimental evolution to test this by quantifying SBGE and sex-specific splicing in 15 *Drosophila melanogaster* populations that evolved for 104 generations in mating treatments that removed mate competition via enforced monogamy, or allowed mate competition in either small, simple or larger, structurally more complex mating environments. Consistent with sex-specific selection affecting SBGE, initially sex-biased genes diverged in expression more among treatments than unbiased genes, and there was greater expression divergence for male-than female-biased genes. It has been suggested the transcriptome should be ‘feminized’ under monogamy because of the removal of sexual selection on males; we did not observe this, likely because selection differs in additional ways between monogamy vs. polygamy. Significant divergence in average expression dimorphism between treatments was observed and, in some treatment comparisons, the direction of the divergence differed across different sex-bias categories. There was not a generalized reduction in expression dimorphism under enforced monogamy.

## Introduction

In many animals, substantial differences between the sexes exist in a myriad of phenotypes involving morphology, behaviour, physiology, and life history. Dimorphism in these phenotypes arises, at least in part, from sex differences in expression of the underlying genes, which themselves are another set of phenotypes for which sexual dimorphism can be considered. Studies across many taxa report the existence of sex-biased expression in a large fraction of genes for any given tissue and/or developmental stage (Grath & Parsch, 2016). However, the extent of transcriptional dimorphism can vary considerably between species (Ingleby et al., 2015), echoing patterns of variation in phenotypic dimorphism.

Contrasting selection between the sexes is widely believed to be the major reason for sexual dimorphism. Sex-specific optima exist for many traits, causing intralocus sexual conflict over optimal expression of underlying genes that are shared between males and females (Bonduriansky & Chenoweth 2009). Such conflict can be resolved via the evolution of sex-biased gene expression (Parsch & Ellegren, 2013). The different reproductive strategies of males and females are thought to be a major source of such sex-specific selection. In particular, selection arising from mate competition, including pre- and post-copulatory sexual selection, often differs between the sexes, shaping the traits that mediate intra- and intersexual interactions. We use “mating system” to refer to the set of inter- and intrasexual interactions related to mating and reproduction.

Mating systems vary widely among species and, hence, this variation is presumed to be a major source of variation in sexual dimorphism, though this has received limited direct attention (Fernandes Martins et al., 2017). At the expression level, Harrison et al. (2015) used a comparative approach to examine the effect of mating system on variation in dimorphism among six bird species. The proportion of genes that were male-biased was positively correlated with presumed indices of both pre- and post-copulatory sexual selection (e.g., sexual ornamentation, sperm number, residual testis mass). Further, the rate of turnover of male-biased genes was positively associated with male sexual ornamentation. This suggests that sexual selection drives the evolution of expression dimorphism. Further evidence consistent with the importance of mating system comes from studies showing that sex-biased gene expression differs between alternative reproductive morphs, for example between ‘dominant’ vs. ‘auxiliary’ males (Dean et al., 2017; Pointer et al., 2013; Stuglik et al., 2014).

Experimental evolution offers a powerful means to directly test whether a change in mating system drives evolutionary changes in the transcriptome. In one such study, Hollis et al. (2014) subjected replicate populations of the naturally polygamous *D. melanogaster* to experimental evolution under two mating systems: polygamy and enforced monogamy (i.e., randomly assigned male-female pairings). The motivation for imposing monogamy was to eliminate sexual selection on males, which is thought to be the primary reason why selection differs between the sexes. Hollis et al. predicted that in the absence of sexual selection on males, a population would no longer be constrained by conflicting selection between the sexes, and hence phenotypes in males and females would evolve towards female optima. Assuming existing patterns of sex-biased gene expression represent only a partial resolution of intralocus sexual conflict, they predicted that evolution under monogamy would result in ‘feminisation’ of the transcriptome (i.e., upregulation of female-biased genes and down-regulation of male-biased genes) in both sexes. These predictions were borne out, with expression being feminised in males and females under enforced monogamy compared to polygamy. However, a similar evolution experiment using *D. pseuodoobscura* found results that were very different from these predictions, with expression being largely masculinised in populations evolving under monogamy compared to polygamy (Veltsos et al., 2017).

Though Hollis et al. (2014) used enforced monogamy with the goal of eliminating sexual selection on males, such manipulations of mating systems are likely to have additional consequences on selection more broadly in both sexes (Rowe & Rundle, 2021). Males are known to inflict harm on females, for example through persistent courtship and toxic seminal fluid proteins (Fowler and Partridge 1989, Partridge and Fowler 1990; Chapman et al. 1995, Arnqvist and Rowe 2005). Under enforced monogamy, there should be strong selection on males to be less harmful to females, and experimental evidence supports this (Holland and Rice 1999; Yun et al. 2021). Selection on females also likely differs between these mating systems. Reduced harm by males may subsequently yield selection against costly female traits involved in avoiding or reducing male harm (Wigby & Chapman, 2004). In addition, under some polygamous but not monogamous conditions, high-quality females may suffer a ‘cost of attractiveness’ as a result of being the targets of preferential male harassment (Long et al., 2009; Yun et al. 2017, MacPherson et al., 2018), ultimately weakening natural selection through females. A change from polygamy to enforced monogamy is thus likely to alter selection in both sexes in a variety of ways, such that contrasts between these mating systems are more nuanced than simply the presence vs. absence of sexual selection. Expression changes may therefore be more difficult to predict than Hollis et al. (2014) suggested. Further complicating this contrast is that ‘polygamy’ can take many forms, characterized by differences in inter- and intrasexual interactions that occur under different contexts (e.g., when reproductive interactions and mating happen in different environments).

Here, we analyze sex-biased gene expression in replicate populations of *D. melanogaster* evolved in three treatments that varied with respect to mating system. One treatment involved the absence of mate competition (MCabs) via the application of enforced monogamy. The remaining two treatments allowed for mate competition, but in distinct ‘mating environments’. In the first of these treatments (MCsim), mate competition occurred in a relatively simple environment (e.g., *Drosophila* vials), similar to the ‘polygamy’ treatment of earlier studies (i.e., Hollis et al. 2014; Veltsos et al. 2017). In the third treatment (MCcom) mate competition occurred in larger, less dense containers with multiple food sources and greater spatial complexity, presumably allowing females to more readily evade males. Consistent with this, in the ‘complex’ relative to the ‘simple’ mating environment, intersexual interactions and mating are less frequent, males are less harmful to females, and females no longer suffer a ‘cost of attractiveness’ that weakens viability selection on them in the simple environment (Yun et al. 2017, 2019; MacPherson et al. 2018). Populations maintained in the complex mating environment also evolved males that are less harmful to females compared to their counterparts from the simple mating environment (Yun et al. 2021), yet these males are highly successful in siring offspring in competition with other males (Yun et al 2019).

Here we analyze gene expression divergence among these three mating treatments from several perspectives. We begin by asking whether divergence among mating treatments is random across the transcriptome with respect to pre-existing sex-bias in expression. If sex-biased gene expression (SBGE) evolves because of differential selection arising from mating interactions, then one would expect that sex-biased genes would be more likely to diverge among mating treatments than would unbiased genes. Second, we contrast female-vs. male-biased genes. Past studies have documented that male-biased genes diverge in expression more rapidly among populations or species than female-biased genes (e.g., Meiklejohn et al. 2003, Zhang et al. 2007, Allen et al. 2018), but this is yet to be considered in relation to variation in mating system. We ask whether male-biased genes are more likely to diverge than female-biased genes in response to changes in mating system.

Third, we examine the directionality of expression changes from the perspective of Hollis et al. (2014), testing their prediction that enforced monogamy should lead to a feminization of the transcriptome relative to polygamous mating systems in which there is a much greater opportunity for sexual selection on males. We compare our enforced monogamy treatment with each of two different polygamous mating treatments in which mate competition occurs in different environments. We also compare the latter two mate competition treatments with one another, asking whether they differ with respect to transcriptome feminization or masculinization.

Changes in dimorphism can occur because of changes in just one sex or both. Moreover, examination of dimorphic traits in each sex separately can provide clues as to whether dimorphism is hindered by intersexual genetic covariances (Lande 1980; Prasad et al. 2007). For such reasons, we examine gene expression in each sex separately, as have past studies (Hollis et al. 2014; Veltsos et al. 2017). For example, Hollis et al. (2014) reported feminization in the transcriptomes of females, and also of males, under enforced monogamy. However, reporting results in this way alone does not provide a clear view of how dimorphism itself changed (this was not a goal of past work). For example, feminization in both sexes under enforced monogamy could mean that dimorphism had increased, decreased, or remained constant, depending on the relative magnitudes of the changes within each sex. Though we are interested in the evolutionary divergence of dimorphic traits within each sex, like past studies (Hollis et al. 2014; Veltsos et al. 2017), we are also interested in whether mating system affects dimorphism per se, so we also explicitly compare dimorphism across mating treatments.

We examine the above questions with respect to expression dimorphism in whole bodies as well as heads. In being less sexually dimorphic, heads offer a point of contrast. Finally, in addition to examining SBGE, we also examine some of these questions with respect to another form of expression dimorphism, sex-specific splicing (SSS), i.e., quantitative differences between the sexes in the relative usage of different isoforms.

## Methods

### Samples for RNAseq

We used 15 populations from a previously described evolution experiment (Yun et al. 2018, 2019, 2021) that were created in September 2014 by sampling from a single laboratory stock population of *D. melanogaster*. During experimental evolution, all populations used here were raised under the same larval conditions consisting of standard cornmeal medium supplemented with 5% NaCl (6% after generation 6) and a constant exposure to 28 C (rather than the standard 25 C of the ancestor) during larval development. The 15 populations were divided equally among three treatments that manipulated the mating system that adults experienced, including the opportunity for mate competition and the environment in which this occurred. The treatments were: mate competition absent (MCabs) via randomly assigned single-pair monogamy, and two polygamy treatments in which mate competition was permitted either in small, structurally simple *Drosophila* vials (MCsim) or in larger, 1.65-L cylindrical containers with added structural complexity (MCcom).

Populations were maintained via 3-week non-overlapping generations. Within a given population, 140 male and 140 female adults spent 6 days each generation in their respective mating treatment. In MCabs, this involved 140 male-female pairs separately allocated to individual wide-mouth straws, while in the polygamy treatments this involved four groups of 35 males and 35 females, each group being held in either separate vials (MCsim) or separate containers (MCcom). Within each of the MCcom containers were five small cups of food (with plastic barriers inserted into the food that further subdivided the surface) and two coiled pipe cleaners, anchored in the lid, that extended into the interior space. After 6 d in these mating treatments, males were discarded and 105 of the surviving females were randomly allocated among seven vials to lay eggs for ∼24 h. Females were subsequently discarded and the resulting offspring developed under the same larval conditions in all populations (i.e., cornmeal food with added salt at 28 C). Adults that eclosed were used to create the next generation’s mating treatments.

After 104 generations of selection, samples for RNAseq were obtained as follows. Flies were reared under a controlled density of 40 larvae per vial on a benign yeast-agar food (no salt). Adults emerged eight days after hatching and were collected under light CO_2_ (< 20 seconds) as virgins (within 8 hours of emergence) and then held in single-sex vials at low density (10 flies per vial). Two days later, flies were processed. For whole body samples, flies were transferred under light CO_2_ (< 20 sec) to microcentrifuge tubes (10 flies per tube). A few minutes later (and well after awakening from CO_2_) tubes were flash frozen in liquid nitrogen and then stored at -80 C. For head samples, the same procedure was used except after flash freezing, tubes were vortex shaken for ca. 10 s to separate heads from bodies. Heads were transferred to a new microcentrifuge tube (8-10 heads per tube) and then stored at -80 C. RNA extraction was performed using ThermoFisher PicoPure RNA Isolation Kit. Paired-end (100 bp) sequencing was performed using Illumina NovaSeq S4. For each of the 15 populations, there was a single replicate of each of the four sample types (i.e., body and head samples of each sex).

One of the female replicates from MCsim treatment was found to have elevated expression of Y-linked genes, indicating possible contamination with male tissue. Consequently, we excluded this replicate from all further analyses.

### Differential Expression between Mating Treatments

Differential expression was analysed between each pair of mating treatments, separately for each sex and tissue. Each RNAseq file was aligned to the *D. melanogaster* reference genome (dos Santos et al., 2015) using STAR v2.7 (Dobin et al., 2013) with default parameters. The resulting alignment files were processed with *htseq-count* to obtain gene-level read counts for each sample (Anders et al., 2015). The R package *DESeq2* (Love et al., 2014) was used to estimate differential expression between pairs of mating treatments. For all genes averaging >50 reads across all replicates, we estimated the log_2_ fold change in expression between treatments (henceforth, ‘treatment effect’). For any genes thus tested, *DESeq2* yields an adjusted p-value by applying Benjamini and Hochberg corrections for multiple testing. A gene was designated to have significant differential expression between treatments if the treatment effect was accompanied by an adjusted p-value < 0.1. Such genes are called ‘Treatment Differentially Expressed genes’ or ‘TDE genes’.

### Sex-Biased Gene Expression

In analyses where SBGE is treated as an independent variable, we used an external dataset (Osada et al., 2017) to characterise genes with respect to sex-biased gene expression for our analyses to avoid circularity that could generate spurious associations if the same data contribute to both dependent and independent variables. This choice for an external dataset is not critical for our purposes; though expression in this external dataset undoubtedly differs somewhat from our own, variation among genes in SBGE tends to be much larger than variation in SBGE for a gene across studies (e.g., genes that are male- or female-biased in one study are generally male- or female-biased in other studies, though they may vary quantitively the magnitude of bias). Even across species separated by millions of years, among-gene SBGE is strongly correlated (Zhang et al. 2007).

The Osada et al. RNAseq dataset consisted of data for whole bodies and heads for a male and female sample from each of 18 lines. Gene-level read counts were obtained as described above. Sex-biased gene expression was estimated using the R package *DESeq2* (Love et al., 2014). In estimating differential expression between sexes, we excluded all genes which averaged <50 reads across all male and female replicates in the given tissue. *DESeq2* yields log_2_ estimates of fold change in male to female expression (“log_2_ FC male/female”). For some analyses, we assigned genes into one of three sex bias categories: female-biased (-∞ < log2FC ≤ -0.5), unbiased (-0.5 < log2FC ≤ 0.5), and male-biased (0.5 < log2FC < ∞).

### Differential Isoform Usage Analysis

In addition to analysing changes to total gene expression, we also considered changes in patterns of alternative splicing using the R package *JunctionSeq* (Hartley and Mullikin 2016). *JunctionSeq* utilises read count data from *QoRTs* (Hartley & Mullikin, 2015), which partitions genes into exons and splice junctions. It then counts the number of reads that overlap an annotated feature (i.e., exon or splice junction). Then, through *JunctionSeq*, generalised linear models (GLM) are used to test for differential splicing. At the gene level, corrections for multiple testing are applied through the Benjamini and Hochberg method, yielding an adjusted p-value for each gene.

We used *QoRTs* to gather read count data from alignment files, previously obtained using *STAR* aligner on the RNA-seq data. Following this, we utilised *JunctionSeq* to analyse differential isoform usage between the sexes, as well as between treatments. In both, we restricted our analyses to features which averaged >50 reads across all replicates. A gene was considered to have significant differential isoform usage between sexes if the gene-wise p-adjusted was below 0.1. For comparisons between treatments, a gene was considered to have significant differential isoform usage if p-adjusted < 0.1. The latter set of genes are referred to as ‘TDS genes’ (for ‘Treatment Differentially Spliced genes’).

### Identification of Gonad-Specific Genes

Patterns of differential expression or isoform usage in whole bodies might be driven in part by the gonads. To examine this possibility, we repeated our analyses for the whole bodies without genes that are expressed specifically in the gonads (‘gonad-specific genes’, GSGs). We identified GSGs using tissue-specific estimates of gene expression from the supplementary data of Witt et al. (2017), which in turn is based on RNAseq data from FlyAtlas2 (Leader et al., 2018). Each gene’s expression in a tissue was expressed in terms of FPKM. We estimated a ‘gonad specificity index’ (GSI) separately for males and females, as per the following expression:

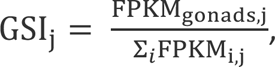

where *j* = sex and *i* = tissue. For the denominator, we only included FPKMs from non-overlapping tissues. Any gene that yielded a GSI > 0.95 in either sex was designated as gonad-specific. We then proceeded to repeat our analyses excluding this set of genes.

## Results

### Divergence in expression is non-random with respect to SBGE

We first examined the frequency of SBGE within each mating treatment. The frequency of genes with significant sex bias in expression was similar across treatments in whole bodies and in heads (Table 1). Though we found statistically significant variation in the frequency of SBGE among treatments in the whole body, these differences are not drastic. This is not surprising because, on a short microevolutionary time, one would not expect large scale expression divergence that would cause genes to change categories of SBGE (especially given that SBGE is fairly consistent across species; Zhang et al. 2007). However, this does not preclude the possibility of many subtle, quantitative changes in expression.

**Table 1:** Frequency of genes with significant sex-biased gene expression (p-adj < 0.05) and significant sex-specific splicing (SSS; p-adj < 0.1) for whole bodies and heads. P-values are from Fisher’s exact tests for non-random association between the percentage of genes with SBGE/SSS and mating treatments. For sex-biased gene expression, the numerator includes only genes that are sex-biased (i.e., |log_2_FC| > 0.5) with p-adj < 0.05, while the denominator includes all genes.

| Sex-biased gene expression (SBGE) |  |  |  |  |  | Sex-specific splicing (SSS) |  |  |  |  |
| --- | --- | --- | --- | --- | --- | --- | --- | --- | --- | --- |
| Tissue | # of genes | % with SBGE |  |  | p-value | # of genes | % with SSS |  |  | p-value |
|  |  | MCabs | MCsim | MCcom |  |  | MCabs | MCsim | MCcom |  |
| Body | 11,717 | 88.4% | 87.6% | 87.2% | $3.4 \times 10^{-5}$ | 7,250 | 49.8% | 48.1% | 49.0% | 0.097 |
| Head | 9,356 | 9.8% | 10.4% | 10.5% | 0.56 | 7,547 | 4.2% | 13.6% | 10.4% | $<2.2 \times 10^{-16}$ |

To that end, we performed pairwise contrasts of mating treatments within each sex to identify genes that were differentially expressed between treatments (hereafter ‘Treatment Differentially Expressed’, TDE, genes). Though a relatively small number of genes qualified as TDE (Table S1), we considered whether these TDE genes are disproportionally represented among different categories of sex-biased expression. If genes that are sex-biased had previously evolved to be so because of a history sex-differential selection arising from mate competition, one might expect that sex-biased genes would be disproportionately likely to diverge among populations evolving in varying mating treatments. However, we may not expect male-vs. female-biased genes to be equally likely to diverge in expression among mating treatments, given results of past studies: relative to the female-biased and unbiased genes, male-biased genes have been reported to have higher rates of evolution in sequence (Zhang et al., 2004) and expression (Meiklejohn et al., 2003).

In female bodies, TDE genes are significantly more common among sex-biased than unbiased genes (Table 2), with the exception of the MCabs-MCsim comparison for which too few TDE genes exist to draw any conclusions. No significant difference is observed in the frequency of TDE genes among unbiased vs. sex-biased genes in comparisons for male bodies, however. Comparing male-vs. female-biased genes, TDE genes are significantly more frequent among male-biased genes in both male and female body samples (again ignoring the MCabs - MCsim comparison due to the low total number of TDE genes). Both of these patterns hold to some degree in heads too: in those comparisons with statistically significant differences, TDE genes are more common among sex-biased than unbiased genes and TDE genes are more common among male-than female-biased genes.

**Table 2:** Results of Fisher’s exact tests for the proportion of treatment differentially expressed (TDE) genes in each sex bias category. Values in parentheses represent the fraction of TDE genes out of all genes in the sex bias category. Genes were assigned to sex-bias categories based on external data source.

| Comparison | Percent TDE genes |  | p-value<br>(UB vs SB) | Percent TDE genes |  | p-value<br>(MB vs FB) |
| --- | --- | --- | --- | --- | --- | --- |
|  | UB | SB |  | FB | MB |  |
| A) Female body |  |  |  |  |  |  |
| MCabs vs MCsim | 0%<br>(0/1693) | 0.03%<br>(2/7025) | 1 | 0.03%<br>(1/3721) | 0.03%<br>(1/3304) | 1 |
| MCabs vs MCcom | 0.5%<br>(9/1693) | 2.0%<br>(137/7025) | **7.0 x 10<sup>-6</sup>** | 0.7%<br>(26/3721) | 3.4%<br>(111/3304) | **2.3 x 10<sup>-16</sup>** |
| MCsim vs McCom | 1.2%<br>(21/1693) | 2.5%<br>(177/7025) | **1.0 x 10<sup>-3</sup>** | 0.8%<br>(29/3721) | 4.5%<br>(148/3304) | **<2.2 x 10<sup>-16</sup>** |
| B) Male body |  |  |  |  |  |  |
| MCabs vs MCsim | 0.5%<br>(9/1699) | 0.25%<br>(23/9101) | 0.082 | 0.1%<br>(5/3546) | 0.3%<br>(18/5555) | 0.13 |
| MCabs vs MCcom | 1.6%<br>(27/1699) | 1.5%<br>(139/9101) | 0.83 | 0.6%<br>(21/3546) | 2.1%<br>(118/5555) | **1.1 x 10<sup>-9</sup>** |
| MCsim vs MCcom | 0.7%<br>(12/1699) | 0.8%<br>(75/9101) | 0.77 | 0.3%<br>(9/3546) | 1.2%<br>(66/5555) | **3.0 x 10<sup>-7</sup>** |
| C) Female head |  |  |  |  |  |  |
| MCabs vs MCsim | 0.7%<br>(54/7767) | 3.7%<br>(32/872) | **1.5 x 10<sup>-11</sup>** | 3.4%<br>(24/698) | 4.6%<br>(8/174) | 0.50 |
| MCabs vs MCcom | 0.3%<br>(27/7767) | 0.8%<br>(7/872) | 0.076 | 0.4%<br>(3/698) | 2.3%<br>(4/174) | **0.032** |
| MCsim vs MCcom | 0.3%<br>(26/7767) | 1.0%<br>(9/872) | **6.6 x 10<sup>-3</sup>** | 0.9%<br>(6/698) | 1.7%<br>(3/174) | 0.39 |
| D) Male head |  |  |  |  |  |  |
| MCabs vs MCsim | 0.5%<br>(35/7774) | 1.1%<br>(10/874) | **0.021** | 1.2%<br>(8/688) | 1.1%<br>(2/186) | 1 |
| MCabs vs MCcom | 1.4%<br>(108/7774) | 3.0%<br>(26/874) | **1.2 x 10<sup>-3</sup>** | 2.2%<br>(15/688) | 5.9%<br>(11/186) | **0.013** |
| MCsim vs MCcom | 0.7%<br>(56/7774) | 1.9%<br>(17/874) | **1.2 x 10<sup>-3</sup>** | 1.7%<br>(12/688) | 2.7%<br>(5/186) | 0.38 |

A possible proximate mechanism for the body results is a change in the relative sizes of gonads (Mank, 2017). We repeated the analysis presented in Table 2 after excluding genes with highly gonad-specific expression. The results remain similar (Table S2). While this does not rule out the possibility that changes in gonads play an important role in expression divergence among treatments, the fact that the patterns are not weakened after removal of gonad-specific genes—along with the existence of the patterns in heads—suggests that there is more to the underlying mechanisms behind expression divergence than solely a change in gonad size.

We analysed the TDE genes for significant enriched biological processes or functions using a gene ontology enrichment test (Eden et al., 2009), but failed to find any significant associations. However, among the 574 TDE genes in total, we found 16 genes that putatively code for seminal fluid proteins (SFPs), identified in a recent study of the seminal fluid proteome (Wigby et al., 2020). One of these 16 putative SFP genes (CG42807) was divergently expressed in two pairwise comparisons (MCabs vs MCcom, MCsim vs MCcom).

### Divergence cannot be summarized as feminization/masculinization of the transcriptome

In the previous section we tested whether different types of genes (with respect to sex-bias) are more likely to diverge, but we did not test for patterns in the direction and magnitude of divergence. In their study, Hollis et al. (2014) predicted that, relative to polygamy, expression would be feminized in populations evolving in monogamy, i.e., female-biased genes evolve to be upregulated in monogamy, while male-biased genes become downregulated. Similar patterns are predicted in both female and male tissues due to the presumed shared genetic architecture that constrains dimorphism under any given mate competition regime (Prasad et al. 2007; Hollis et al. 2014).

We evaluated expression divergence in our populations from this Hollis et al. perspective by examining divergence separately for female- and male-biased genes, focusing on the contrasts of our monogamy treatment (MCabs) with each the two polygamy treatments (MCsim, MCcom). To visualise and test for patterns of transcriptomic feminization/masculinization, we used the ‘treatment effect’ for each gene as a measure of expression change, defined as the logarithmic ratio of expression in treatment 1 relative to treatment 2. A positive treatment effect in a male-biased gene, for instance, would imply that its expression is relatively masculinized in treatment 1; for female-biased genes, a positive treatment effect would similarly imply relatively feminized expression in treatment 1. (We included unbiased genes for completeness but these are not pertinent to the Hollis et al. prediction.) Results (Fig. 1) suggest that the relationship between treatment effect and sex bias is inconsistent across treatment pairs, sexes and sample types (head/body). Matching the Hollis et al. (2014) prediction, there is an overall feminization of the female transcriptome in MCabs relative to the other two treatments, with a significant increase in expression of female-biased genes and a significant decrease in expression of male-biased genes in MCabs (Fig. 1A-B). In male bodies (Fig. 1D-E), the net changes are smaller, but more importantly, the pattern is different. Though female-biased genes have increased expression in MCabs relative to the other two treatments (similar to the female body result), male-biased genes are not significantly reduced in expression in MCabs relative to MCsim and, in fact, have significantly increased expression in MCabs relative to MCcom (i.e., opposite the prediction). A further layer of complication is added when examining the heads, where the results differ between female and male samples and in neither sex is there a consistent pattern of the predicted feminization of MCabs (Fig 1G-H, K-L).

**Figure 1:**
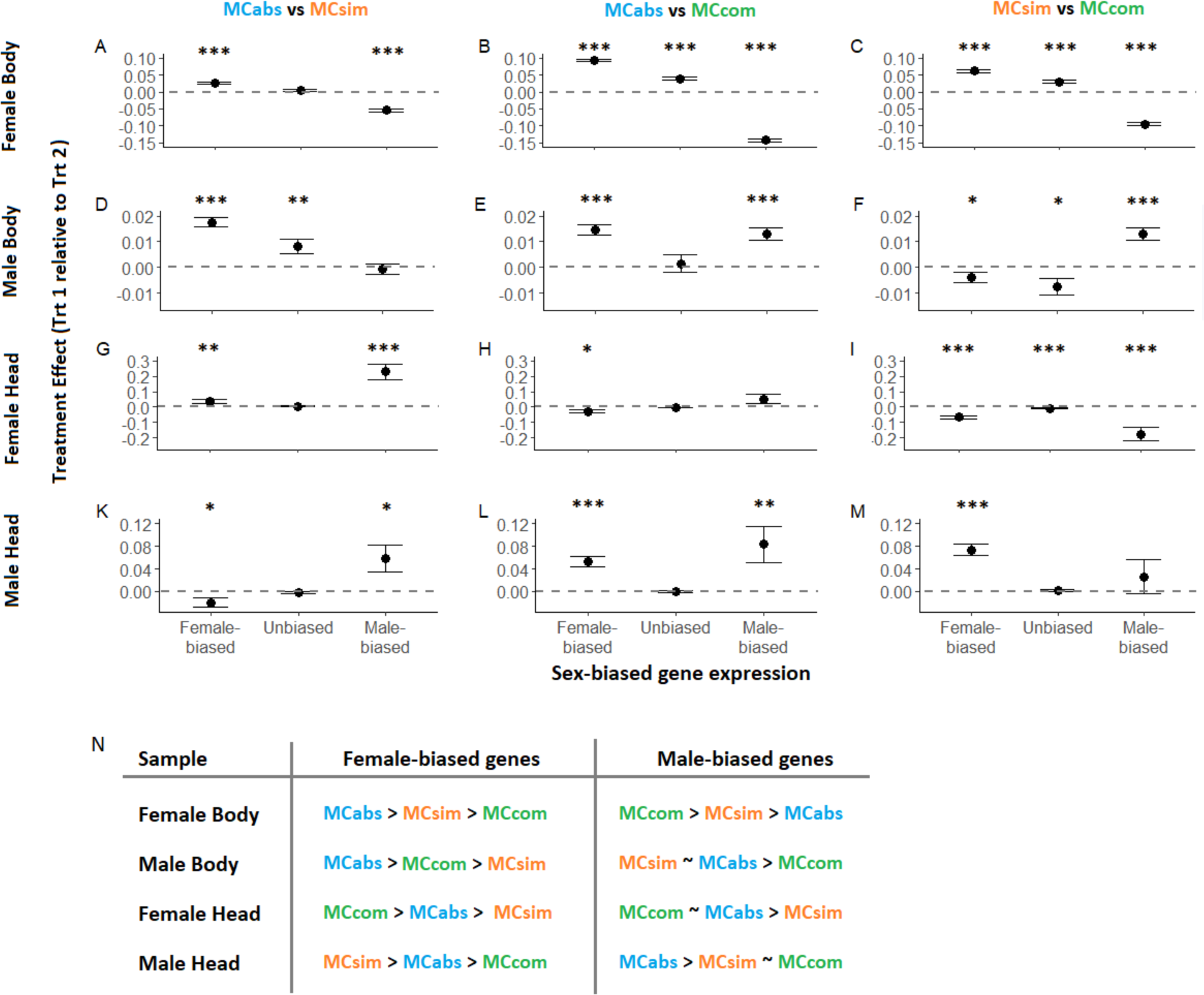
Average treatment effect (± 1 SE) for three sex bias categories for female body (A-C), male body (D-F), female head (G-I), male head (K-M), in pairwise comparisons between mating treatments. Asterisks denote the mean treatment effect is significantly different from zero, determined using one-sample permutation tests (i.e., by changing the sign of the treatment effect of a randomly chosen 50% of the genes and re-calculating the mean in each permutation). Significance is denoted as follows: * p < 0.05, ** p < 0.01, *** p < 0.001. Panel N summarises the order of expression levels for female- and male-biased genes in panels A-M; tilda (∼) denotes no significant difference. An analogous figure from an analysis excluding gonad-specific genes is shown in Fig. S1; most of the patterns are very similar. Genes were assigned to sex-bias categories based on external data source.

Fig. 1 depicts mean treatment effects in discrete, and somewhat arbitrarily-bounded categories of sex bias. We also performed LOESS regressions of treatment effect against continuously varying sex bias to discern any patterns missed by treating variation in sex-biased gene expression as discrete. These regressions (Fig. S2) show that the relationship between treatment effect and sex bias can vary within a sex bias category, a nuance not captured in the discrete version of the plots. However, the overall sign of mean treatment effect for a sex bias category is largely consistent in the two analyses.

As another approach to evaluating the Hollis et al. prediction, we looked for evidence of net changes in feminization/masculinization via a different method using only the TDE genes. In female bodies, there is a significant net feminization of TDE genes in MCabs relative to MCcom (Fig. S3), similar to the results depicted in Fig. 1 and matching the Hollis et al. prediction. In male bodies, there is a significant net masculinization of TDE genes in MCabs relative to MCcom (Fig. S3), opposite to the Hollis et al. prediction. In most other contrasts, there is no significant net feminization or masculinization.

### Changes in degree of sexual dimorphism

In the previous section we examined expression changes for each sex separately, but these within-sex changes are not directly informative of changes in sexual dimorphism. Here we investigate differences between treatments in the degree of expression dimorphism across genes with different levels of sex-bias as determined from an external dataset (see Methods). We used LOESS regressions to evaluate how treatment differences in ‘local average dimorphism’ varied with sex-bias (where local average refers to the difference in dimorphism averaged across a local range of sex-bias). Patterns were complex (Fig. 2) though treatment differences in local average dimorphism did tend to be larger for sex-biased than unbiased genes. Among the sex-biased genes, the magnitude and even direction of treatment differences in local average dimorphism varied between, but also within, sex-bias categories. In bodies, local average dimorphism is highest in MCsim for both highly female- and male-biased genes, but patterns differ for moderately-biased genes (summarized in Fig. 2G). In heads, where phenotypic sexual dimorphism is not as marked, changes in local average dimorphism are detectable despite the relatively low number of sex-biased genes. Female-biased genes are, on average, most dimorphic in MCcom, but male-biased genes are least dimorphic in MCcom. Analyses excluding gonad-specific genes yielded nearly identical results.

**Figure 2:**
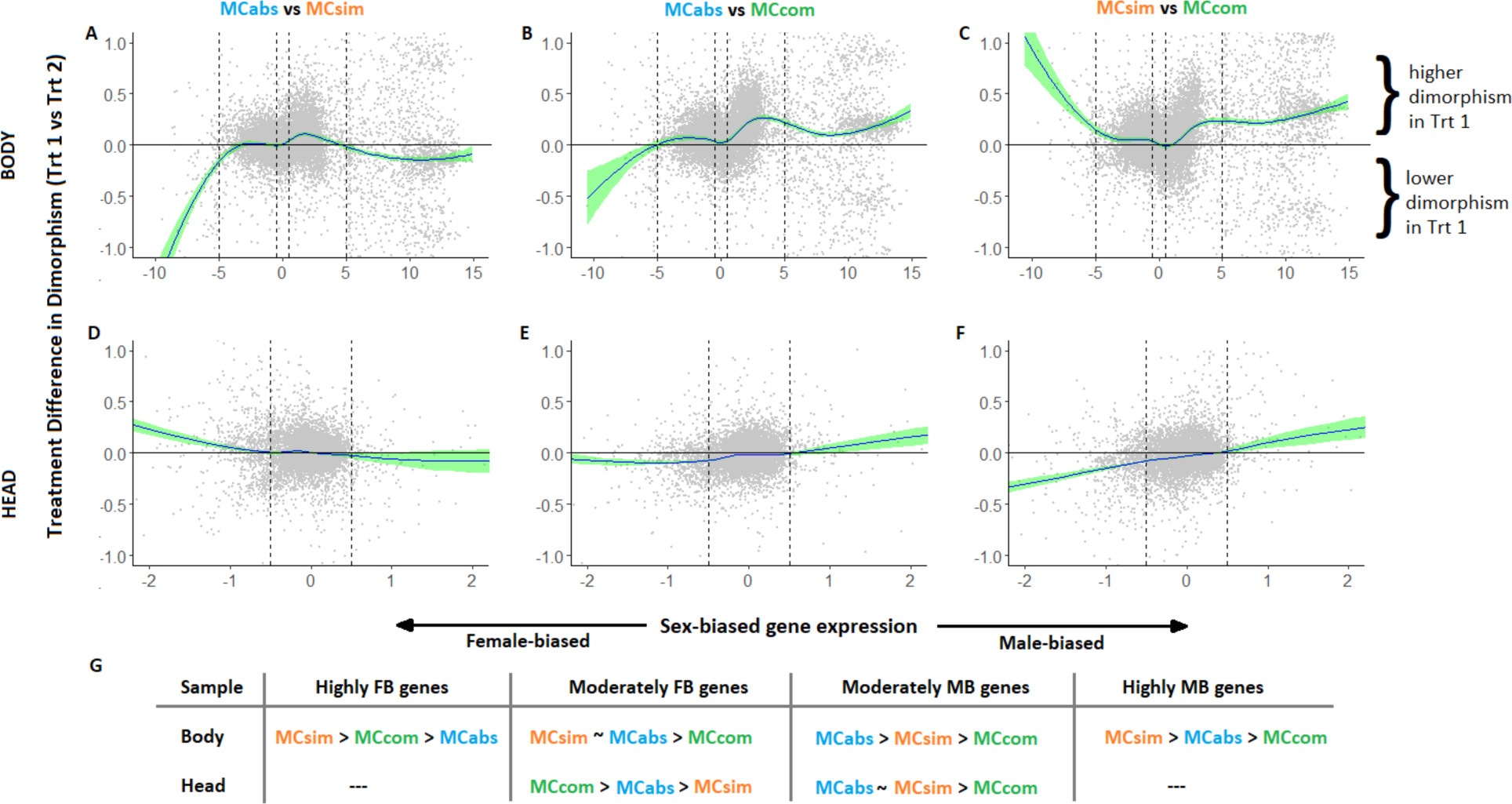
Treatment difference in dimorphism as a function of sex-biased gene expression in A-C) body and D-F) head. The vertical axis is (sign of sex bias)*(difference in dimorphism between treatments) so that positive (negative) values indicate increased (reduced) dimorphism in treatment 1 relative to treatment 2. In panels A-F, the fitted line is obtained from LOESS regressions (fit using *geom_smooth* in the R package *ggplot2* with the width of the sliding-window set at 0.5); 95% confidence intervals are indicated in green. The plots show a subset of the genes within the vertical axis range [-1,1] to better display the pattern for the majority of genes. Dashed vertical lines demarcate the sex bias categories listed in the panel G. Panel G summarises the relative strength of dimorphism among treatments for each SBGE category. Sex-bias values were determined from an external data source. The categories used in panel G are designated as follows: highly FB genes = (-∞,-5], moderately FB genes = (-5, -0.5], moderately MB genes = [0.5, 5), [5, ∞). No highly male- or female-biased genes are depicted for heads due to the low numbers of such genes.

### Differential splicing amongst mating treatments

We also examined sex-specific splicing (SSS), testing whether alternative splicing differs between mating treatments. Congruent to our analysis for TDE genes, we tested for genes with significant differential splicing between treatments, i.e., ‘treatment differentially spliced’ (TDS) genes. Frequencies of TDS genes are in the range of 0.2-2.3% of all genes analysed (Table S3). We performed Fisher’s exact tests to ascertain if TDS genes are disproportionately represented among sex-biased genes. In contrast to the analogous test for TDE genes (Table 2, S2), there was no over-representation of TDS genes among sex-biased genes, though there was some evidence of TDS genes being more common among male-than female-biased genes (Tables S4, S5). Considering TDE and TDS together, there was significant overlap between these in most sample types (Table S6).

We also tested for sex-specific splicing (SSS) within each treatment. The proportion of SSS genes was similar among treatments, with one exception: the proportion of such genes is noticeably lower in the heads in MCabs than the other two treatments (Table 1). The lower frequency of SSS in MCabs heads is unlikely to be due to reduced power to detect SSS: MCabs heads had the highest mean coverage (Table S7) and there was no comparable in reduction in the frequency of sex-biased expression (Table 1). This difference in treatment effects on the prevalence SBGE and SSS suggests that these two forms of expression dimorphism evolve somewhat independently.

## Discussion

Differential selection on the sexes is thought to drive the evolution of sexual dimorphism in gene expression (Parsch & Ellegren, 2013). Selection arising from sexual interactions and mate competition is hypothesized to be a major cause of this differential selection. Thus, variation in mate competition among populations or species should be a major source of variation in SBGE across such taxonomic groups. Here, we tested this idea by measuring shifts in gene expression in response to evolutionary manipulations of mate competition in *D. melanogaster*. The MCabs treatment eliminates mate competition through enforced monogamy, while the two other treatments (MCsim, MCcom) incorporate mate competition in differing mating environments that we know alter intersexual interactions and the selection these generate (MacPherson et al., 2018; Yun et al., 2017, 2019).

An implicit assumption in evaluating the predictions raised in the Introduction is that the observed differences in expression are, at least to some extent, adaptive. This warrants consideration. The *DEseq2* analysis identifies changes that are consistent across populations (i.e., evolutionarily independent replicates) within selective treatments; that is, these are expression differences that can be attributed to the effect of the selective treatments. Previous studies of fitness-related traits (Yun et al. 2019; 2021) show that these populations have adapted to their mating treatments. However, as with other “evolve and resequence” (E&R) expression studies (e.g., Huang et al. 2014, Hollis et al. 2014, Veltsos et al. 2017, Hsu et al. 2020), we do not have direct evidence that the observed expression differences contribute to the observed fitness changes. Though we suspect the expression changes will tend to be adaptive, some of the expression changes are likely incidental outcomes of linked selection, which can cause parallel, non-adaptive changes in E&R studies.

Differences in the mating regimes could also result in differences in the strength of drift among treatments. However, drift is not expected to create consistent changes in allele frequencies across evolutionarily independent replicates, and thus would not typically contribute to a statistically-identified “selective treatment effect”. There is one potentially important exception. A treatment with stronger drift would have populations carrying a heavier burden of deleterious alleles and thereby lower average fitness within each replicate, though the causative alleles would differ among replicate populations. If the effects of deleterious alleles at different sites are somewhat interchangeable in that they contribute to a common latent variable such as “condition” (*sensu* Rowe and Houle 1994), which itself affects expression, then differences in drift would contribute to a “treatment effect”. Indeed, Wyman et al. (2010) showed that rearing flies on a poor quality diet, which could be viewed as a means to reduce “condition”, altered sex-biased gene expression. However, this is a specific manipulation of the environment and not a demonstration that different sets of deleterious alleles would cause a consistent expression effect. There is some evidence that lowering diet quality is not representative of the effects of deleterious alleles (Bonduriansky et al. 2015).

Perhaps most importantly, Singh and Agrawal (2022) estimated *N_e_*/*N* in *D. melanogaster* from direct measures of male and female variance in fitness under three mating treatments that closely parallel those used here, and found similar values for all three (see also Pischedda et al. 2015.). The common intuition that mate competition will necessarily and dramatically alter *N_e_* can be misleading for both theoretical reasons (see Crow and Morton 1955; Singh and Agrawal 2021) and an empirical one: the variance in female fitness increases under enforced monogamy (Long et al. 2009; Singh and Agrawal 2021). In summary, we have reasons to believe that *N_e_* is similar among our treatments; it is nevertheless possible that there are subtle differences in the strength of drift and these could contribute to the treatment-level expression differences reported here, but only under the specific circumstances outlined above. Given we know that these populations have evolved adaptively with respect to their mating treatments (Yun et al. 2019; 2021), we suspect adaptive evolution is an important process behind the observed expression changes. Nonetheless, as we interpret our results below, it is important to bear in mind that some of the observed expression changes are likely non-adaptive, resulting from linked selection or drift. (Similar issues would apply to related studies, e.g., Hollis et al (2014), Veltsos et al. (2017).)

If a history of sex-differential selection arising from mate competition is responsible for pre-existing SBGE, then we expect that selection on genes that are initially sex-biased is more likely to be sensitive to a change in mate competition than selection on unbiased genes. Thus, we predicted that sex-biased genes would be more likely to diverge among mate competition treatments than unbiased genes. The results largely supported this prediction (Table 2); all seven of the comparisons in which there was a significant difference were in the predicted direction (though not all comparisons yielded a significant difference).

Though the patterns are consistent with the prediction, other explanations likely also contribute to this pattern. Core house-keeping genes will tend to be unbiased and their expression may be under strong stabilizing selection that changes little across any of a large range of environments. For this reason, we might predict that unbiased genes are less likely to diverge in expression than sex-biased genes across any environmental change, not just changes to mate competition. A better test, which may be possible via meta-analysis in future work, would be to ask whether the preferential divergence of sex-biased relative to unbiased genes is more pronounced in response to mate competition treatments than it is with other types of environmental changes.

We also compared the relative frequency of DE genes in male-versus female-biased genes. Past studies have shown that male-biased genes evolve more rapidly than female-biased genes, both in DNA sequence (Meisel, 2011; Zhang et al., 2004) and gene expression (Allen et al., 2018; Meiklejohn et al., 2003), though some studies describe more mixed results (Whittle & Johannesson, 2013; Yang et al., 2016). Adaptive evolution in response to sexual selection has been invoked as an explanation (Ellegren & Parsch, 2007), with some studies suggesting that male-biased genes are targeted more often by strong positive selection (Pröschel et al., 2006; Zhang et al., 2004; but see Singh and Agrawal 2023) presumably resulting from reproductive interactions. Our manipulation of mate competition resulted in more divergence in expression of male-than female-biased genes (Table 2): in the six comparisons where there was a significant difference, male-biased genes had a higher propensity for expression divergence than female-biased genes. In some sense, this matches with fitness data showing stronger signatures of local adaptation to the mate competition environment in males than females (Yun et al. 2019), suggesting greater phenotypic divergence in males than females. However, Table 2 contrasts male- and female-biased genes within each sex, rather than contrasting divergence of males versus that of females. There are not obvious differences in the proportion of significantly diverged genes between male and female samples. This apparent discrepancy could be reconciled if the expression divergence of male-biased genes has greater phenotypic and fitness consequences in males than females.

Hollis et al. (2014) predicted that enforced monogamy would result in a “feminization” of the transcriptome (i.e., increased expression of female-biased genes and decreased expression of male-biased genes) in both sexes. This prediction has now been tested in three experiments with highly heterogeneous results. The data of Hollis et al. (2014) matched their prediction, while the results of Veltsos et al. (2017) were mixed but largely in the opposite direction. Our results are mixed with respect to the predicted changes.

The basis of the Hollis et al. prediction is that the primary consequence of enforced monogamy is the removal of sexual selection on males. However, enforced monogamy not only removes sexual selection, but it imposes (potentially strong) selection on males to inflict less harm on their mates (Holland and Rice 1999; Martin and Hosken 2003; Crudginton et al. 2005, 2010; Yun et al. 2021). This can lead to changes in selection on female resistance to harm (Holland & Rice 1999, Martin and Hosken 2003). Even in the absence of evolved changes in male harm, there are strong reasons to suspect that enforced monogamy will change selection on females because males will be unable to bias their attention towards particular phenotypes (Long et al., 2009; Arbuthnott and Rundle 2012; Chenoweth et al. 2015; Yun et al. 2017). Considering this multitude of possible changes in selection, it is difficult to predict how the transcriptome will respond, and it seems unlikely that net change will be consistent or easily classifiable as ‘feminization’ or ‘masculinization.’

Because selection in enforced monogamy can differ from selection under mate competition in a variety of ways, other details could become important to how divergence occurs. These three studies differ in various ways which could, in principle, contribute to the among-study heterogeneity in results (e.g., species studied, nature of starting variation (e.g., Hollis et al. (2014) add new mutations via chemical mutagenisis to standing variation), food, temperature, density and population size, and maintenance procedures (Li Richter & Hollis 2021)). One potentially important aspect is the precise nature of mate competition, which can take many forms (e.g., there is no one true form of “polygamy” for a “monogamy vs. polygamy” contrast). Our study directly shows that differences in how mate competition occurs has major consequences for expression evolution. Within the context of our experiment, where most other factors are not varied among treatments, the transcriptional differences between our two different mate competition treatments (i.e., MCsim vs. MCcom) are generally as large as those between either one of these with the enforced monogamy treatment (MCabs).

The hypothesis that the removal of sexual selection would ‘feminize’ the transcriptome was tested in a very different way by Parker et al. (2019). They examined expression changes in five asexual species of stick insects relative to their sexual sister species, under the idea that asexual females do not share a gene pool with sexually-selected males. Contrary to expectation, Parker et al. (2019) found evidence that asexual females had evolved ‘masculinized’ expression. They suggested that the most plausible explanation was that much of feminized expression patterns in sexual females arises via traits involved with female-male interactions and, because asexual females do not mate or interact with males, expression evolves to be less ‘feminized’. By this explanation, asexual species are not an ideal counterfactual for inferring how sharing a gene pool with sexually-selected males constrains the evolution of expression in sexual females because asexual females also experience substantially different selection than sexual females.

Though selection arising from mate competition in sexual species is almost certainly an important driver of SBGE (Parsch & Ellegren, 2013), there is little direct evidence (Harrison et al. 2014). It seems intuitive that, in the absence of mate competition, expression dimorphism would become reduced. For example, monogamous bird species are often less dimorphic with respect to plumage (and other traits) than non-monogamous species, and presumably this reflects a reduction in expression dimorphism for genes affecting such traits. Yet, it is unclear what the prediction for genome-wide expression dimorphism should be as intralocus conflict on expression can easily exist under monogamy given that males and females play different roles in reproduction and offspring rearing, even if overt traits such as plumage do not experience divergent selection.

While the underlying motivation for the studies of Hollis et al. (2014) and Veltsos et al. (2017) was based around sex-biased gene expression, neither attempted to examine how enforced monogamy changed expression dimorphism per se. We examined how the average treatment effect on expression dimorphism varied across local ranges of sex bias as determined in LOESS regressions. We found differences in ‘local average dimorphism’ among mating regimes. These changes in local average dimorphism tend to occur to a greater extent among sex-biased than unbiased genes, and more so in bodies, which have much more sex-biased gene expression than heads (Fig. 2). However, we did not find a consistent reduction in local average dimorphism between the enforced monogamy treatment and the two treatments with mate competition; while some categories of sex-biased genes became less dimorphic under monogamy, others became more so (Fig. 2). For example, relative to MCsim, MCabs evolved reduced local average dimorphism of strongly sex-biased genes but increased dimorphism of moderately male-biased genes in body samples. However, one aspect of expression was consistent with the intuition of reduced dimorphism under monogamy: there was a notable reduction in the frequency of genes with sex-specific splicing in heads of MCabs relative to either of the other two treatments, though not in bodies (Table 1). While our experiment is the result of more than 100 generations of selection, this is a relatively short period relative to other evolutionary time scales (e.g., sister species); perhaps a more consistent reduction in expression dimorphism would be evident after enough time. Nonetheless, this study shows that, though expression dimorphism readily evolves among mate competition environments, enforced monogamy does not necessarily cause a rapid and consistent reduction in dimorphism.

The most striking change in local average dimorphism occurred between the two treatments involving mate competition, with MCsim showing greater dimorphism in bodies than MCcom across all sex bias categories (Fig. 2C). Relative to MCcom, the MCsim environment is one in which intersexual interactions, including mating, occur more frequently (Yun et al. 2017, 2019); exposure to males is more harmful to females in the simple than complex environment and MCsim males have evolved to be more harmful (Yun et al. 2017, 2021). Thus, it is reasonable to infer that interlocus sexual conflict is relatively more important in MCsim than MCcom. However, this does not provide an obvious explanation for the observed difference in expression dimorphism, as dimorphism is thought to evolve in response to intra-, rather than interlocus, conflict.

In this study, we have treated each gene’s expression as a trait (i.e., a measurable property) and examined patterns in how these ‘traits’ change in response to an evolutionary manipulation of mate competition. Just as with traditional traits (i.e., morphological, behavioral, or physiological phenotypes), multiple possible genetic mechanisms could underly each change and a single genetic change could affect multiple traits. At one (unlikely) extreme, every gene expression change could be due to a change in that gene’s *cis*-regulatory region. At the other extreme, a single genetic change could affect expression levels of many genes, possibly by changing the size of different organs or relative abundance of cell types within organs (Stewart et al. 2010). The truth likely lies somewhere between these two extremes. In an evolutionary manipulation of mating system in *Drosophila pseudoobscura*, Wiberg et al. (2021) found that genomically diverged sites were enriched near sites with expression divergence. That result would not be expected if a very small number of genetic changes underlay most expression differences, but it also does not imply that most expression changes are due to individual *cis*-regulatory changes.

It is not our objective here to resolve this issue for our own experiment (and we have little ability to do so). In the spirit of many past transcriptomic studies, we test for patterns among different ‘types’ of traits (e.g., genes with male-biased, female-biased, or unbiased expression). From this perspective, each gene represents a separate instance of that trait ‘type’ but it is important to remember that variation in each of those individual ‘traits’ need not be governed by independent proximate mechanisms. This unknown level of independence should temper inferences about genetic changes and the true nature and number of targets of selection (e.g., could some patterns result from changes in the relative size of gonads and, if so, why has selection caused such changes?). Our study clearly reveals patterns of expression evolution across SBGE categories in response to changes in mate competition, though we are limited in interpreting the reasons for these patterns. The type of work presented here represents only one step towards understanding the relationship between mate competition and SBGE, which can help evaluate hypotheses in the literature and provide fodder for new ones.

## Author Contributions

PM performed bioinformatic data processing. AA and HR conceived the project. PM, HR, and AA contributed to data analysis and writing of the manuscript.

## Data availability

Upon acceptance, RNAseq data will be submitted to the Sequence Read Archive (NIH) and analysis scripts will be made available via GitHub.

## Conflict of Interest

Authors have no conflicts of interest.

## Acknowledgements

We thank George Sandler for doing the RNA extractions.

## Funding

This work was supported by the Natural Sciences and Engineering Research Council of Canada (Discovery grants to AFA & HDR).

## Notes

### Competing Interest Statement

The authors have declared no competing interest.

### Summary of Updates

Minor updates, primarily to Discussion.

## Literature Cited

Allen, S. L., R. Bonduriansky, and S. F. Chenoweth. 2018. Genetic constraints on microevolutionary divergence of sex-biased gene expression. Philos. Trans. R. Soc. Lond. B Biol. Sci. 373:20170427.

Anders, S., P. T. Pyl, and W. Huber. 2015. HTSeq--a Python framework to work with high-throughput sequencing data. Bioinformatics 31: 166–169.

Arbuthnott, D., and H. D. Rundle. 2012. Sexual selection is ineffectual or inhibits the purging of deleterious mutations in *Drosophila melanogaster*. Evolution 66:2127–2137.

Arnqvist, G., and L. Rowe. 2005. Sexual conflict. Princeton University Press, Princeton, NJ. Bonduriansky, R., and S. F. Chenoweth. 2009. Intralocus sexual conflict. Trends Ecol. Evol. 24:280–288.

Bonduriansky, R., M. A. Mallet, D. Arbuthnott, V. Pawlowsky-Glahn, J. J. Egozcue, and H. D. Rundle. 2015. Differential effects of genetic vs. environmental quality in Drosophila melanogaster suggest multiple forms of condition dependence. Ecol. Lett. 18:317–26.

Chapman, T., L. F. Liddle, J. M. Kalb, M. F. Wolfner, and L. Partridge. 1995. Cost of mating in *Drosophila melanogaster* females is mediated by male accessory gland products. Nature 373:241–244. nature.com.

Chenoweth, S. F., N. C. Appleton, S. L. Allen, and H. D. Rundle. 2015. Genomic evidence that sexual selection impedes adaptation to a novel environment. Curr. Biol. 25:1860–1866.

Crow J. F., and N. E. Morton NE. 1955. Measurement of gene frequency drift in small populations. Evol. 9:202–14.

Crudgington, H. S., A. P. Beckerman, L. Brüstle, K. Green, and R. R. Snook. 2005. Experimental removal and elevation of sexual selection: does sexual selection generate manipulative males and resistant females? Am. Nat. 165: S72–87.

Crudgington, H. S., S. Fellows, and R. R. Snook. 2010. Increased opportunity for sexual conflict promotes harmful males with elevated courtship frequencies. J. Evol. Biol. 23:440–446.

Dean, R., A. E. Wright, S. E. Marsh-Rollo, B. M. Nugent, S. H. Alonzo, and J. E. Mank. 2017. Sperm competition shapes gene expression and sequence evolution in the ocellated wrasse. Mol. Ecol. 26:505–518.

Dobin, A., C. A. Davis, F. Schlesinger, J. Drenkow, C. Zaleski, S. Jha, P. Batut, M. Chaisson, and T. R. Gingeras. 2013. STAR: ultrafast universal RNA-seq aligner. Bioinformatics 29:15–21.

dos Santos, G., A. J. Schroeder, J. L. Goodman, V. B. Strelets, M. A. Crosby, J. Thurmond, D. B. Emmert, W. M. Gelbart, and FlyBase Consortium. 2015. FlyBase: introduction of the *Drosophila melanogaster* Release 6 reference genome assembly and large-scale migration of genome annotations. Nucleic Acids Res. 43:D690–7.

Eden, E., R. Navon, I. Steinfeld, D. Lipson, and Z. Yakhini. 2009. GOrilla: a tool for discovery and visualization of enriched GO terms in ranked gene lists. BMC Bioinformatics 10:48.

Ellegren, H., and J. Parsch. 2007. The evolution of sex-biased genes and sex-biased gene expression. Nat. Rev. Genet. 8:689–698.

Fernandes Martins, M. J., G. Hunt, R. Lockwood, J. P. Swaddle, and D. J. Horne. 2017. Correlation between investment in sexual traits and valve sexual dimorphism in Cyprideis species (Ostracoda). PLoS One 12:e0177791.

Fowler, K., and L. Partridge. 1989. A cost of mating in female fruitflies. Nature 338:760–761.

Grath, S., and J. Parsch. 2016. Sex-biased gene expression. Annu. Rev. Genet. 50:29–44.

Harrison, P. W., A. E. Wright, F. Zimmer, R. Dean, S. H. Montgomery, M. A. Pointer, and J. E. Mank. 2015. Sexual selection drives evolution and rapid turnover of male gene expression. Proc. Natl. Acad. Sci. U. S. A. 112:4393–4398.

Hartley, S. W., and J. C. Mullikin. 2016. Detection and visualization of differential splicing in RNA-Seq data with JunctionSeq. Nucleic Acids Res. 44:e127.

Holland, B., and W. R. Rice. 1999. Experimental removal of sexual selection reverses intersexual antagonistic coevolution and removes a reproductive load. Proc. Natl. Acad. Sci. U. S. A. 96:5083– 5088.

Hollis, B., D. Houle, Z. Yan, T. J. Kawecki, and L. Keller. 2014. Evolution under monogamy feminizes gene expression in *Drosophila melanogaster*. Nat. Commun. 5:3482.

Ingleby, F. C., I. Flis, and E. H. Morrow. 2014. Sex-biased gene expression and sexual conflict throughout development. Cold Spring Harb. Perspect. Biol. 7:a017632.

Lande, R. 1980. Sexual dimorphism, sexual selection, and adaptation in polygenic characters. Evolution 34:292–305. JSTOR.

Leader, D. P., S. A. Krause, A. Pandit, S. A. Davies, and J. A. T. Dow. 2018. FlyAtlas 2: a new version of the *Drosophila melanogaster* expression atlas with RNA-Seq, miRNA-Seq and sex-specific data. Nucleic Acids Res. 46:D809–D815.

Li Richter, X.-Y., and B. Hollis. 2021. Softness of selection and mating system interact to shape trait evolution under sexual conflict. Evolution 75:2335–2347.

Long, T. A. F., A. Pischedda, A. D. Stewart, and W. R. Rice. 2009. A cost of sexual attractiveness to high-fitness females. PLoS Biol. 7:e1000254.

Love, M. I., W. Huber, and S. Anders. 2014. Moderated estimation of fold change and dispersion for RNA-seq data with DESeq2. Genome Biol. 15:550.

MacPherson, A., L. Yun, T. S. Barrera, A. F. Agrawal, and H. D. Rundle. 2018. The effects of male harm vary with female quality and environmental complexity in *Drosophila melanogaster*. Biol. Lett. 14:20180443.

Martin, O. Y., and D. J. Hosken. 2003. Costs and benefits of evolving under experimentally enforced polyandry or monogamy. Evolution 57:2765–2772.

Meiklejohn, C. D., J. Parsch, J. M. Ranz, and D. L. Hartl. 2003. Rapid evolution of male-biased gene expression in *Drosophila*. PNAS 100:9894–9899.

Osada, N., R. Miyagi, and A. Takahashi. 2017. Cis- and trans-regulatory effects on gene expression in a natural population of *Drosophila melanogaster*. Genetics 206:2139–2148.

Parker D.J., J. Bast, K. Jalvingh, Z. Dumas, M. Robinson-Rechavi, and T. Schwander. 2019. Sex-biased gene expression is repeatedly masculinized in asexual females. Nat. Comm. 10:4638.

Parsch, J., and H. Ellegren. 2013. The evolutionary causes and consequences of sex-biased gene expression. Nat. Rev. Genet. 14:83–87.

Partridge, L., and K. Fowler. 1990. Non-mating costs of exposure to males in female *Drosophila melanogaster*. J. Insect Physiol. 36:419–425.

Pointer, M. A., P. W. Harrison, A. E. Wright, and J. E. Mank. 2013. Masculinization of gene expression is associated with exaggeration of male sexual dimorphism. PLoS Genet. 9:e1003697.

Prasad, N. G., S. Bedhomme, T. Day, and A. K. Chippindale. 2007. An evolutionary cost of separate genders revealed by male-limited evolution. Am. Nat. 169:29–37.

Pröschel, M., Z. Zhang, and J. Parsch. 2006. Widespread adaptive evolution of *Drosophila* genes with sex-biased expression. Genetics 174:893–900.

Rowe, L., and H. D. Rundle. 2021. The alignment of natural and sexual selection. Annu. Rev. Ecol. Evol. Syst. 52:499–517.

Singh, A., and A. F. Agrawal. 2022. Sex-specific variance in fitness and the efficacy of selection. Am. Nat. 199: 587–602.

Singh, A., and A. F. Agrawal. 2023. Two forms of sexual dimorphism in gene expression in *Drosophila melanogaster*: their coincidence and evolutionary genetics. Mol. Biol. Evol. 40: msad091.

Stewart, A. D., A. Pischedda, and W. R. Rice. 2010. Resolving intralocus sexual conflict: genetic mechanisms and time frame. J. Hered. 101:S94–9.

Stuglik, M. T., W. Babik, Z. Prokop, and J. Radwan. 2014. Alternative reproductive tactics and sex-biased gene expression: the study of the bulb mite transcriptome. Ecol. Evol. 4:623–632.

Veltsos, P., Y. Fang, A. R. Cossins, R. R. Snook, and M. G. Ritchie. 2017. Mating system manipulation and the evolution of sex-biased gene expression in *Drosophila*. Nat. Commun. 8:2072.

Whittle, C. A., and H. Johannesson. 2013. Evolutionary dynamics of sex-biased genes in a hermaphrodite fungus. Mol. Biol. Evol. 30:2435–2446.

Wiberg, R. A. W., P. Veltsos, R. R. Snook, and M. G. Ritchie. 2021. Experimental evolution supports signatures of sexual selection in genomic divergence. Evol Lett 5:214–229.

Wigby, S., N. C. Brown, S. E. Allen, S. Misra, J. L. Sitnik, I. Sepil, A. G. Clark, and M. F. Wolfner. 2020. The Drosophila seminal proteome and its role in postcopulatory sexual selection. Philos. Trans. R. Soc. Lond. B Biol. Sci. 375:20200072.

Wigby, S., and T. Chapman. 2004. Female resistance to male harm evolves in response to manipulation of sexual conflict. Evolution 58:1028–1037.

Witt, E., N. Svetec, S. Benjamin, and L. Zhao. 2021. Transcription factors drive opposite relationships between gene age and tissue specificity in male and female *Drosophila* gonads. Mol. Biol. Evol. 38:2104–2115.

Yang, L., Z. Zhang, and S. He. 2016. Both male-biased and female-biased genes evolve faster in fish genomes. Genome Biol. Evol. 8:3433–3445.

Yun, L., A. F. Agrawal, and H. D. Rundle. 2021. On male harm: How it is measured and how it evolves in different environments. Am. Nat. 198:219–231.

Yun, L., M. Bayoumi, S. Yang, P. J. Chen, H. D. Rundle, and A. F. Agrawal. 2019. Testing for local adaptation in adult male and female fitness among populations evolved under different mate competition regimes. Evolution 73:1604–1616.

Yun, L., P. J. Chen, K. E. Kwok, C. S. Angell, H. D. Rundle, and A. F. Agrawal. 2018. Competition for mates and the improvement of nonsexual fitness. Proc. Natl. Acad. Sci. U. S. A. 29:201805435.

Yun, L., P. J. Chen, A. Singh, A. F. Agrawal, and H. D. Rundle. 2017. The physical environment mediates male harm and its effect on selection in females. Proc. Biol. Sci. 284:20170424.

Zhang, Y., D. Sturgill, M. Parisi, S. Kumar, and B. Oliver. 2007. Constraint and turnover in sex-biased gene expression in the genus *Drosophila*. Nature 450:233–237.

Zhang, Z., T. M. Hambuch, and J. Parsch. 2004. Molecular evolution of sex-biased genes in *Drosophila*. Mol. Biol. Evol. 21:2130–2139.

