## Supplementary Material for "The evolution of sexual dimorphism in gene expression in response to a manipulation of mate competition" (from another version of this preprint: version 1, 2023-08-17)

Table S1. Percentage of significant (p-adjusted < 0.1) treatment differentially expressed (TDE) genes detected for each treatment pair in males and females. The number of TDE genes are shown in parentheses.

| **Sample** | **Comparison** | **Females** | | **Males** | |
|---|---|---|---|---|---|
| | | Total # of genes | % of TDE genes | Total # of genes | % of TDE genes |
| Body | MCabs vs MCsim | 9199 | 0.03% (N = 3) | 12163 | 0.32% (N = 39) |
| | MCabs vs MCcom | 9178 | 1.69% (N = 155) | 12165 | 1.59% (N = 194) |
| | MCsim vs MCcom | 9205 | 2.28% (N = 210) | 12149 | 0.87% (N = 106) |
| Head | MCabs vs MCsim | 9615 | 1.08% (N = 104) | 9647 | 0.60% (N = 58) |
| | MCabs vs MCcom | 9446 | 0.58% (N = 55) | 9611 | 1.70% (N = 163) |
| | MCsim vs MCcom | 9486 | 0.58% (N = 55) | 9589 | 0.99% (N = 95) |

Table S2. Results of Fisher's exact tests for the proportion of treatment differentially expressed (TDE) genes in each sex bias category, excluding gonad-specific genes. Values in parentheses represent the number of TDE genes out of the total number of genes in the sex bias category.

| Sample | Comparison | Percentage of TDE genes | | p-value (UB vs SB) | Percentage of TDE genes | | p-value (MB vs FB) |
|---|---|---|---|---|---|---|---|
| | | UB | SB | | FB | MB | |
| Female body | MCabs vs MCsim | 0% (0/1691) | 0.03% (2/6987) | 1 | 0.03% (1/3712) | 0.03% (1/3275) | 1 |
| | MCabs vs MCcom | 0.5% (9/1691) | 1.9% (136/6985) | **$1.0 \times 10^{-5}$** | 0.7% (26/3712) | 3.4% (110/3275) | **$2 \times 10^{-16}$** |
| | MCsim vs MCcom | 1.2% (21/1691) | 2.5% (177/6985) | 0.053 | 0.8% (29/3712) | 4.5% (148/3275) | **$2 \times 10^{-16}$** |
| Male body | MCabs vs MCsim | 0.5% (9/1696) | 0.3% (22/7808) | 0.10 | 0.1% (5/3540) | 0.4% (17/4268) | 0.051 |
| | MCabs vs MCcom | 1.6% (27/1696) | 1.7% (130/7808) | 0.91 | 0.6% (21/3540) | 2.6% (109/4268) | **$1.7 \times 10^{-12}$** |
| | MCsim vs MCcom | 0.7% (12/1696) | 0.9% (69/7808) | 0.56 | 0.3% (9/3540) | 1.4% (60/4268) | **$1.3 \times 10^{-8}$** |
| Female head[1] | MCabs vs MCsim | 0.7% (53/7743) | 3.6% (31/868) | **$1.5 \times 10^{-11}$** | 3.3% (23/695) | 4.6% (8/173) | 0.50 |
| | MCabs vs MCcom | 0.3% (27/7743) | 0.8% (7/868) | 0.076 | 0.4% (3/695) | 2.3% (4/173) | **0.032** |
| | MCsim vs MCcom | 0.3% (26/7743) | 1.0% (9/868) | **$6.5 \times 10^{-3}$** | 0.9% (6/695) | 1.7% (3/173) | 0.39 |
| Male head[1] | MCabs vs MCsim | 0.5% (35/7749) | 1.1% (10/871) | **0.021** | 1.2% (8/687) | 1.1% (2/184) | 1 |
| | MCabs vs MCcom | 1.4% (106/7749) | 3.0% (26/871) | **$7.1 \times 10^{-4}$** | 2.2% (15/687) | 6.0% (11/184) | **0.013** |
| | MCsim vs MCcom | 0.7% (54/7749) | 2.0% (17/871) | **$5.5 \times 10^{-4}$** | 1.7% (12/687) | 2.7% (5/184) | 0.38 |

[1]Gonad-specific genes were identified using a separate data set using a criterion given in the Methods. A very small number of genes identified as "gonad-specific" had expression in the head so the gene set is here is very slightly different than that in Table 2.

Table S3. Percentage of significant (p-adjusted < 0.1) 'Treatment Differentially Spliced' (TDS) genes for each treatment comparison in males and females. The raw number of TDS genes for a given comparison are shown in parentheses.

| **Sample** | **Comparison** | **Females** | | **Males** | |
|---|---|---|---|---|---|
| | | Total # of genes | % of TDS genes | Total # of genes | % of TDS genes |
| Body | MCabs vs MCsim | 7516 | 0.29% (N = 22) | 9549 | 0.16% (N = 15) |
| | MCabs vs MCcom | 7465 | 1.00% (N = 75) | 9520 | 1.14% (N = 109) |
| | MCsim vs MCcom | 7471 | 2.28% (N = 170) | 9543 | 0.46% (N = 44) |
| Head | MCabs vs MCsim | 7703 | 0.23% (N = 18) | 7715 | 0.40% (N = 31) |
| | MCabs vs MCcom | 7613 | 0.64% (N = 49) | 7704 | 0.53% (N = 41) |
| | MCsim vs MCcom | 7644 | 0.54% (N = 41) | 7696 | 0.81% (N = 62) |

Table S4: Results of Fisher's exact tests for the proportion of treatment differentially spliced (TDS) genes in each sex bias category. Values in parentheses represent the number of TDS genes out of the total number of genes in the sex bias category.

| Sample | Comparison | Percentage of TDS genes | | p-value (UB vs SB) | Percentage of TDS genes | | p-value (MB vs FB) |
|---|---|---|---|---|---|---|---|
| | | UB | SB | | FB | MB | |
| Female body | MCabs vs MCsim | 0.4% (6/1462) | 0.3% (16/5761) | 0.42 | 0.3% (8/3164) | 0.3% (8/2597) | 0.82 |
| | MCabs vs MCcom | 1.7% (25/1462) | 0.9% (50/5761) | **0.008** | 0.6% (19/3164) | 1.2% (31/2597) | **0.02** |
| | MCsim vs MCcom | 2.2% (32/1462) | 2.3% (135/5761) | 0.77 | 2.2% (71/3164) | 2.5% (64/2597) | 0.60 |
| Male body | MCabs vs MCsim | 0.2% (3/1474) | 0.1% (10/7314) | 0.47 | 0.06% (2/2984) | 0.2% (8/4330) | 0.21 |
| | MCabs vs MCcom | 1.5% (22/1474) | 1.1% (84/7314) | 0.29 | 0.7% (22/2984) | 1.4% (62/4330) | **$7.0 \times 10^{-3}$** |
| | MCsim vs MCcom | 0.3% (4/1474) | 0.5% (38/7314) | 0.30 | 0.2% (7/2984) | 0.7% (31/4330) | **$4.5 \times 10^{-3}$** |
| Female head | MCabs vs MCsim | 0.2% (12/6647) | 0.7% (5/715) | **0.02** | 0.8% (5/592) | 0% (0/123) | 0.59 |
| | MCabs vs MCcom | 0.6% (41/6647) | 1% (7/715) | 0.22 | 1.0% (6/592) | 0.8% (1/123) | 1 |
| | MCsim vs MCcom | 0.6% (37/6647) | 0.6% (4/715) | 1 | 0.7% (4/592) | 0% (0/123) | 1 |
| Male head | MCabs vs MCsim | 0.4% (28/6663) | 0.1% (1/715) | 0.36 | 0.2% (1/580) | 0% (0/135) | 1 |
| | MCabs vs MCcom | 0.5% (36/6663) | 0.4% (3/715) | 1 | 0.3% (2/580) | 0.7% (1/135) | 0.47 |
| | MCsim vs MCcom | 0.9% (59/6663) | 0.4% (3/715) | 0.28 | 0.5% (3/580) | 0% (0/135) | 1 |

Table S5. Results of Fisher's exact tests for the proportion of treatment differentially spliced (TDS) genes in each sex bias category, excluding gonad-specific genes. Values in parentheses represent the fraction of TDS genes out of the total number of genes in the sex bias category.

| Sample | Comparison | Percentage of TDS genes | | p-value (UB vs SB) | Percentage of TDS genes | | p-value (MB vs FB) |
|---|---|---|---|---|---|---|---|
| | | UB | SB | | FB | MB | |
| Female body | MCabs vs MCsim | 0.4% (6/1460) | 0.3% (16/5729) | 0.43 | 0.2% (8/3155) | 0.3% (8/2564) | 0.8 |
| | MCabs vs MCcom | 1.7% (25/1460) | 0.9% (50/5729) | **9.0 x $10^{-3}$** | 0.6% (19/3155) | 1.2% (31/2564) | **0.02** |
| | MCsim vs MCcom | 2.2% (32/1460) | 2.4% (135/5729) | 0.77 | 2.2% (71/3155) | 2.5% (64/2564) | 0.60 |
| Male body | MCabs vs MCsim | 0.2% (3/1471) | 0.1% (9/6508) | 0.47 | 0.07% (2/2979) | 0.2% (7/3529) | 0.19 |
| | MCabs vs MCcom | 1.5% (22/1471) | 1.1% (73/6508) | 0.23 | 0.7% (22/2979) | 1.4% (51/3529) | **8.0 x $10^{-3}$** |
| | MCsim vs MCcom | 0.3% (4/1471) | 0.5% (30/6508) | 0.38 | 0.2% (7/2979) | 0.6% (23/3529) | **0.02** |
| Female head | MCabs vs MCsim | 0.2% (12/6638) | 0.7% (5/713) | **0.02** | 0.8% (5/590) | 0% (0/123) | 0.59 |
| | MCabs vs MCcom | 0.6% (41/6638) | 1% (7/713) | 0.22 | 1.0% (6/590) | 0.8% (1/123) | 1 |
| | MCsim vs MCcom | 0.6% (37/6638) | 0.6% (4/713) | 1 | 0.7% (4/590) | 0% (0/123) | 1 |
| Male head | MCabs vs MCsim | 0.4% (28/6655) | 0.1% (1/713) | 0.36 | 0.2% (1/578) | 0% (0/135) | 1 |
| | MCabs vs MCcom | 0.5% (36/6655) | 0.4% (3/713) | 1 | 0.3% (2/578) | 0.7% (1/135) | 0.47 |
| | MCsim vs MCcom | 0.9% (59/6655) | 0.4% (3/713) | 0.28 | 0.5% (3/578) | 0% (0/135) | 1 |

Table S6. Results of Fisher's exact tests for the overlap in TDE and TDS genes.

| Comparison | Sample | Total number of genes | Number of genes that are TDS and TDE/total number of TDE genes | Number of genes that are TDS and non-TDE/total number of non-TDE genes | p-value |
|---|---|---|---|---|---|
| **MCabs vs MCsim** | Female body | 7416 | 0/(1+0) = 0% | 22/(22+7388) = 0.3% | 1 |
| | Male body | 9358 | 0/(0+1) = 0% | 15/(15+9310) = 0.2% | 1 |
| **MCabs vs MCcom** | Female body | 7362 | 2/(2+1) = 66.7% | 73/(73+7157) = 1.0% | **$3.2 \times 10^{-4}$** |
| | Male body | 9337 | 11/(11+1) = 91.6% | 97/(97+9087) = 1.1% | **$<2.2 \times 10^{-16}$** |
| **MCsim vs MCcom** | Female body | 7373 | 15/(15+1) = 93.8% | 152/(152+7044) = 2.1% | **$<2.2 \times 10^{-16}$** |
| | Male body | 9356 | 5/(5+1) = 83.3% | 39/(39+9238) = 0.4% | **$1.1 \times 10^{-11}$** |
| **MCabs vs MCsim** | Female head | 7591 | 0/(0+1) = 0% | 18/(18+7493) = 0.2% | 1 |
| | Male head | 7601 | 0/(0+1) = 0% | 31/(31+7518) = 0.4% | 1 |
| **MCabs vs MCcom** | Female head | 7512 | 6/(6+1) = 85.7% | 43/(43+7430) = 0.6% | **$4.0 \times 10^{-13}$** |
| | Male head | 7591 | 6/(6+1) = 85.7% | 35/(35+7429) = 0.5% | **$1.3 \times 10^{-13}$** |
| **MCsim vs MCcom** | Female head | 7537 | 3/(3+1) = 75.0% | 37/(37+7460) = 0.5% | **$5.6 \times 10^{-7}$** |
| | Male head | 7578 | 10/(10+1) = 90.9% | 52/(52+7453) = 0.7% | **$<2.2 \times 10^{-16}$** |

Table S7. Mean coverage of genes in each mating treatment

| Treatment | Mean coverage ± SE in body | Mean coverage ± SE in heads |
|---|---|---|
| **MCabs** | 760.92 ± 69.19 | 1012.38 ± 126.99 |
| **MCsim** | 762.23 ± 43.83 | 993.56 ± 93.71 |
| **MCcom** | 714.77 ± 52.60 | 799.75 ± 39.91 |

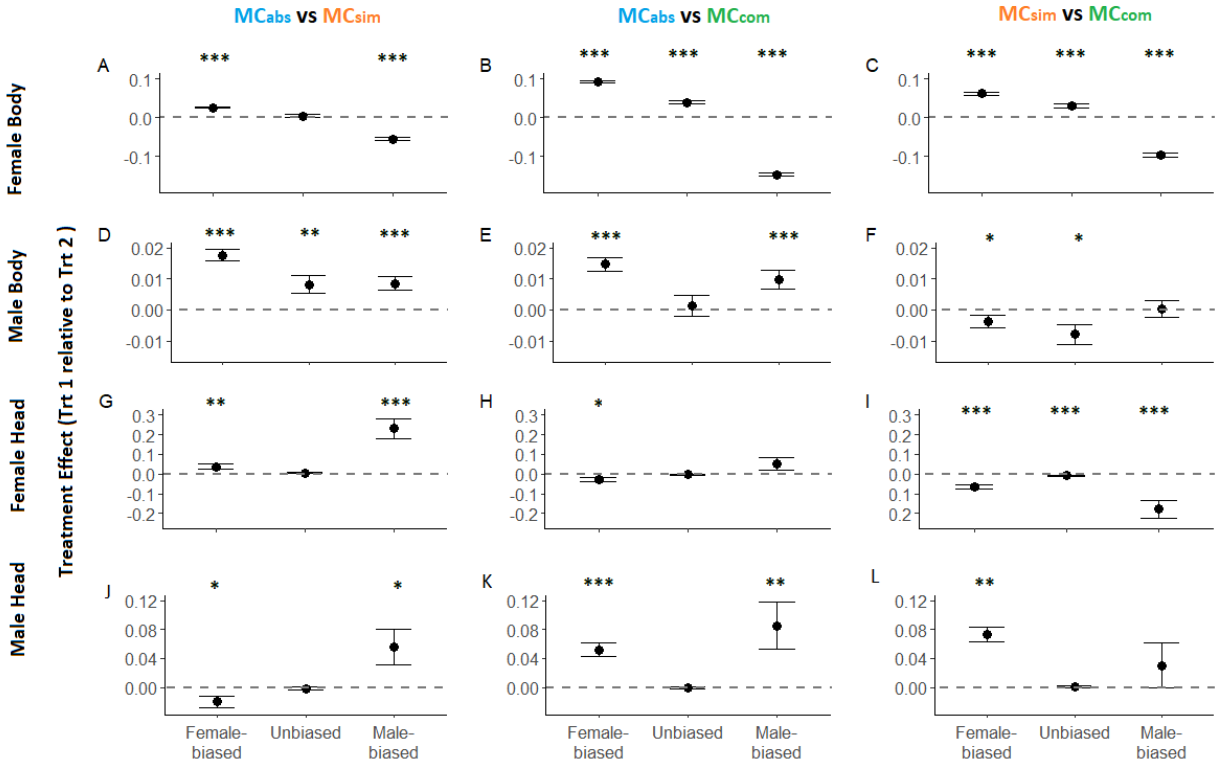


M

| Sample | Female-biased genes | Male-biased genes |
|---|---|---|
| **Female Body** | MCabs > MCsim > MCcom | MCcom > MCsim > MCabs |
| **Male Body** | MCabs > MCcom > MCsim | MCabs > MCsim ~ MCcom |
| **Female Head** | MCcom > MCabs > MCsim | MCcom ~ MCabs > MCsim |
| **Male Head** | MCsim > MCabs > MCcom | MCabs > MCsim ~ MCcom |

Figure S1: Average treatment effect (± 1 SE) in pairwise comparisons between mating treatments for the three sex bias categories for female body (A-C), male body (D-F), female head (G-I), and male head (J-L) after excluding gonad-specific genes. Treatment effect is defined as the $\log_2$ ratio of expression in treatment 1 relative to treatment 2. Asterisks denote the mean treatment effect is significantly different from zero, determined using one-sample permutation tests (i.e., by changing the sign of the treatment effect of a randomly chosen 50% of the genes and re-calculating the mean in each permutation). Significance is denoted as follows: * $p < 0.05$, ** $p < 0.01$, *** $p < 0.001$. Panel N summarizes the order of expression levels for female- and male-biased genes in panels A-M; tilde (~) denotes no significant difference.

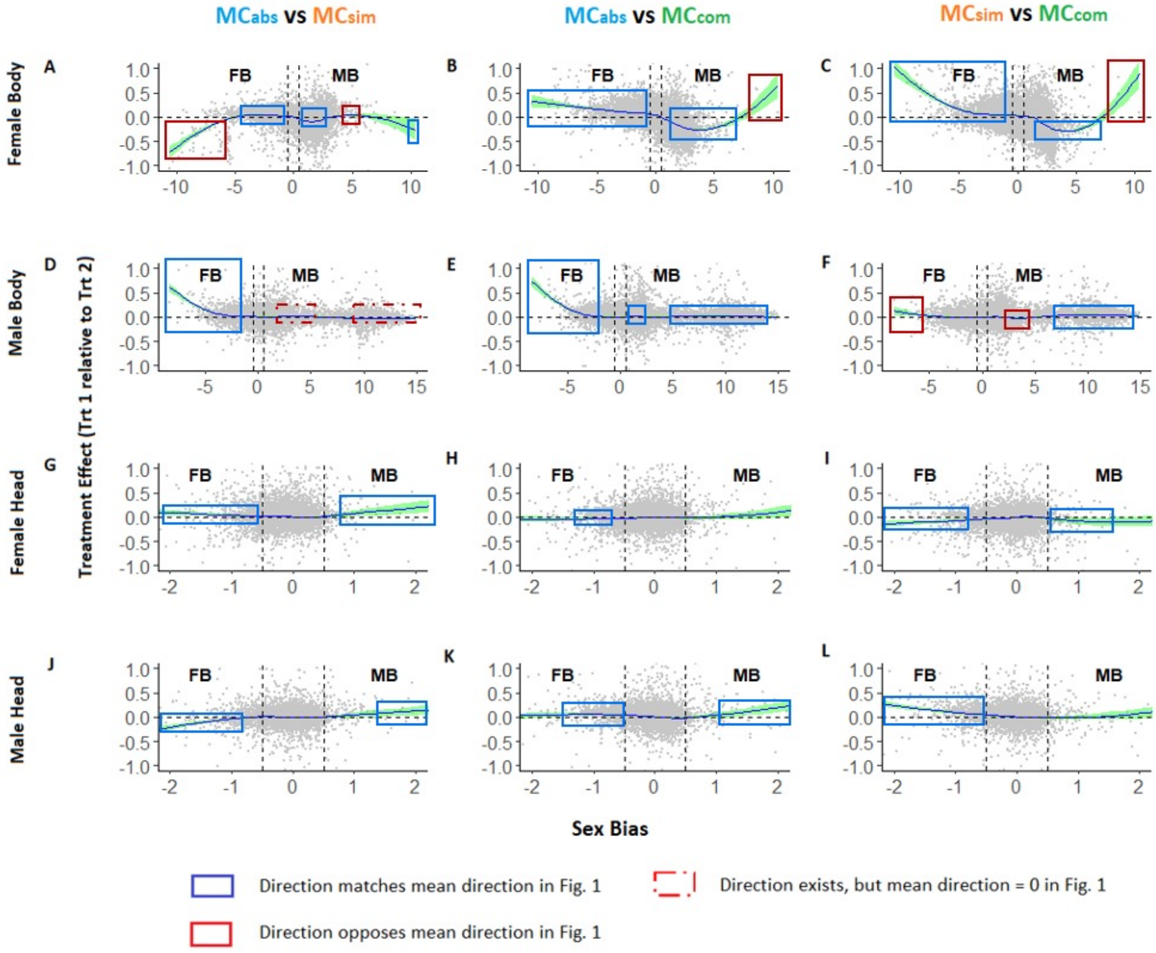


Fig S2. LOESS regressions of treatment effect on sex-biased gene expression (fit using *geom_smooth* in the R package *ggplot2* with the width of the sliding-window set at 0.5) for A-C) female body, D-F) male body, G-I) female head, J-L) male head. The rectangular boxes highlight regions where the 95% confidence intervals (indicated in green) for the LOESS fit do not overlap 0. The direction of the effect relative to that shown in Fig. 1 is indicated by the color of the box (blue = same direction as in Fig. 1; red = opposite to Fig. 1; dashed red = no significant direction exists in Fig. 1). The vertical axis has been zoomed tis only displayed in the range [-1,1] to aid visualization (i.e., some points are not shown but were included in LOESS regression).

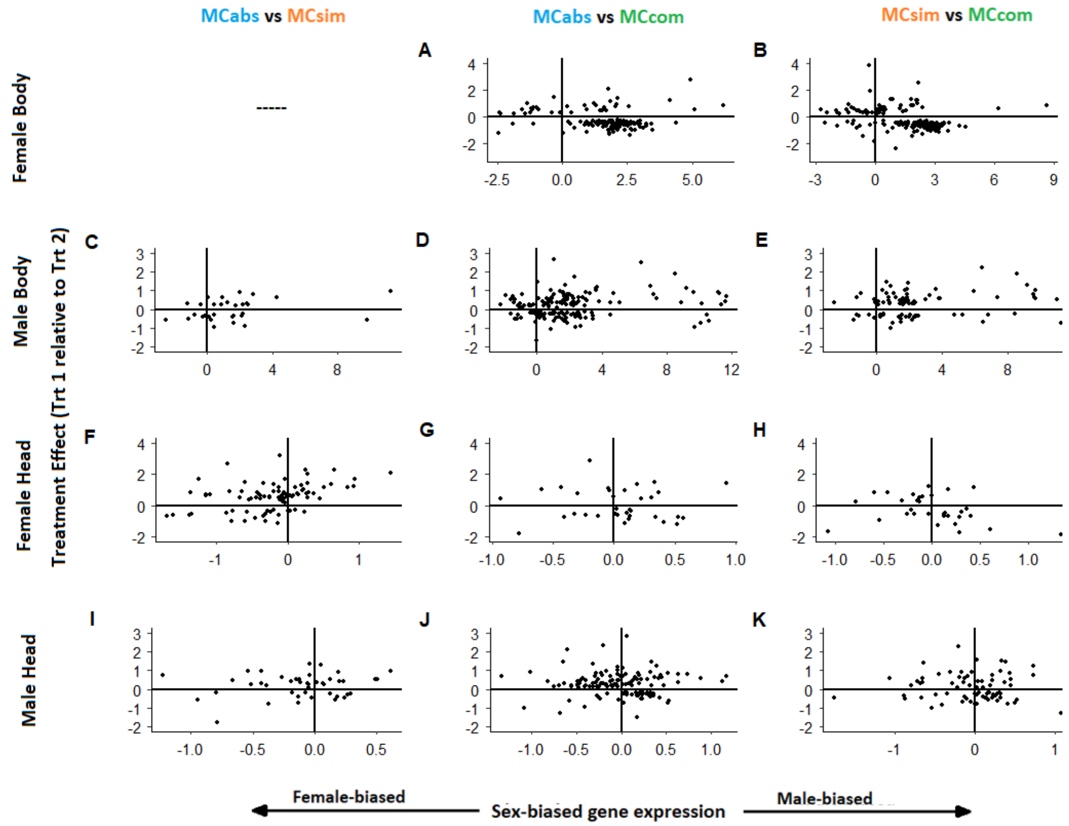


L Summary of expression changes in Treatment 1 relative to Treatment 2

| | MCabs vs MCsim | MCabs vs MCcom | MCsim vs MCcom |
|---|---|---|---|
| Female Body | --- | feminisation *** | feminisation *** |
| Male Body | --- | masculinisation *** | masculinisation *** |
| Female Head | --- | --- | --- |
| Male Head | masculinisation ** | --- | --- |

Figure S3. Treatment effect and sex-biased gene expression for A-B) female body, C-E) male body, F-H) female head, I-K) male head. Plot for MCabs vs MCsim in female body is not shown, as the comparison yielded only two TDE genes. Panel L summarizes the net change in ‘sexualized expression’ in treatment 1 relative to treatment 2. To do so, a relative sexualization score was calculated for each TDE gene as $z$ = TE*SBGE where TE is the treatment effect (i.e., log2 expression of treatment 1 relative to treatment 2) and SBGE is the sex effect (i.e., log2 expression of males relative to females, measured from an external dataset). A positive $z$ values implies a relative change towards ‘masculinization’ in treatment 1 (e.g., decreased expression of a female-biased gene or increased expression of a male-biased gene); a negative value implies a relative change towards ‘feminization’ of treatment 1. To estimate the net change in relative ‘sexualized expression’, we calculated the mean value of $z$ across all TDE genes in each pairwise comparison. To assess significance, we performed one-sample permutation tests with 10,000 iterations (by changing the sign of the treatment effect of a randomly chosen 50% of the genes and re-calculating the mean in each permutation). “---” denotes no significant net change in sexualized expression (or insufficient data in the case of MCabs vs MCsim for female body).
